# Rapid fabrication of collagen bundles mimicking tumor-associated collagen signatures

**DOI:** 10.1101/815662

**Authors:** Xiangyu Gong, Jonathan Kulwatno, K. L. Mills

**Affiliations:** Department of Mechanical, Aerospace, and Nuclear Engineering, Rensselaer Polytechnic Institute, 110 8th St, Troy, NY 12180; Department of Biomedical Engineering, Rensselaer Polytechnic Institute, 110 8th St, Troy, NY 12180; Center for Biotechnology and Interdisciplinary Studies, Rensselaer Polytechnic Institute, 110 8th St, Troy, NY 12180

**Keywords:** Tumor microenvironment, Collagen, Hydrogels, 3D culture, Microfluidics, Invasion

## Abstract

Stromal collagen surrounding a solid tumor tends to present as dense, thick bundles. The collagen bundles are remodeled during tumor progression: first tangential to the tumor boundary (indicating growth) and later perpendicular to the tumor boundary (indicating likely metastasis). Current reconstituted-collagen *in vitro* tumor models are unable to recapitulate the *in vivo* structural features of collagen bundling and alignment. Here, we present a rapid yet simple procedure to fabricate collagen bundles with an average thickness of 9 μm, compared to the reticular dense collagen nanofiber (∼900 nm-diameter, on average) prepared using common protocols. The versatility of the collagen bundles was demonstrated with their incorporation into two *in vitro* models where the thickness and alignment of the collagen bundles resembled the various *in vivo* arrangements. First, collagen bundles aligned by a microfluidic device elicited cancer cell contact guidance and enhanced their directional migration. Second, the presence of the collagen bundles in a bio-inert agarose hydrogel was shown to provide a highway for cancer cell invasion. The unique structural features of the collagen bundles advance the physiological relevance of *in vitro* collagen-based tumor models for accurately capturing cancer cell-stroma interactions.

## 1. Introduction

Tissue architecture, such as that provided by collagen and other fibrous proteins, profoundly influences cell behavior and functionality by providing the resident cells with structural support and mechanical cues^1, 2^. Cancer cells in turn remodel collagen structures during cancer progression via contraction^3, 4^, crosslinking^5, 6^, and degradation^7^. In tumor tissue explants from mice, Provenzano *et al.* observed that collagen at the tumor-stroma interface bundles together and becomes denser at tumor initiation^8, 9^. At progressive stages, the microscale collagen bundles orient tangential to the tumor boundary, and eventually reorient perpendicular to the tumor boundary. The authors termed these three stages of temporal remodeling of the collagen microstructure the tumor-associated collagen signatures (TACS-1, -2, and -3) and intended to define visual hallmarks that characterize critical stages of tumor development. Tumor cell behavior is affected by the collagen architecture at every stage, including increased invasiveness associated with radially aligned collagen^8, 10^. Although these thick bundled collagen strands were commonly associated with tumor progression *in vivo*, *in vitro* models often overlook these distinct architectural features. *In vitro* platforms that accurately capture the key features of TACS are necessary to determine the mechanisms of cancer cell metastasis.

Researchers have sought to recapitulate the biophysical properties of the tumor-associated extracellular matrix (ECM) *in vitro* using biological or synthetic hydrogels^11^, or other polymeric structures, such as polycaprolactone (PCL) fibers^12^. Given that collagen is the most abundant ECM protein^13^, reconstituted Type I collagen hydrogels are popular models, which are characterized by nanoscale collagen fibers physically bound to form an isotropic and porous structure^14^. Cancer cell division^15^, migration^16, 17^, invasive phenotypes^6, 18, 19^ have been observed in collagen gels of varied pore size, stiffness, or crosslink density. But the nanoscale, isotropic collagen fibers are not able to fully recapitulate collagen alignment and the level of bundling of TACS.

Attempts to engineer more physiologically relevant tumor microenvironments are plentiful. Collagen nanofibers have been aligned with an applied magnetic field^20^, mechanical stretching^21^, and microfluidics^22^, although the readily achievable fiber diameter of the collagen (tens to hundreds of nanometers) is not comparable to the microscale fibers observed in the *in vivo* tumor microenvironment. In need of larger (microscale), aligned fibrous structures, polymeric fibers have been created by electrospinning^23^. These fibers were shown to promote a mesenchymal morphology of epithelial cells; however, the synthetic materials used lack ligand-binding sites as collagen does for cell functions^24^ or the proper mechanical properties possessed by native tissue. Moreover, electrospinning requires a special experimental setup that is not accessible in every lab. Recently, polyacrylamide gel-based microgroove topographies coated with collagen have been made to study cancer cell contact guidance and cell spreading on microscale structures^25, 26^. Although the size, spacing, and alignment of these polymer fibers or microgrooves could be systematically produced to mimic those particular features of TACS, the cancer cells were seeded on a surface with topographic features, which lacks the 3D confinement nature of the *in vivo* tissue. Thus, to effectively study cancer cell-stroma interactions *in vitro*, there stands a need for a model made from a native tissue component that better mimics bundling level or alignment of TACS.

In this study, we discovered a simple method to rapidly fabricate microscale bundles of type I collagen, with bundle sizes greater than any that have been previously reported for any *in vitro* collagen-based models. This method may be used to address the gap between current *in vitro* tumor models and *in vivo* tumor stromal architectures. We first introduce the fabrication procedure and its mechanism. The thickness of the collagen bundles was then characterized and compared to collagen fibers prepared by a traditional protocol. We further demonstrated the ability to incorporate microscale collagen bundles into two different *in vitro* systems–a 3D microfluidic device and an agarose-based collagen composite gel (co-gel), where the alignment and thickness of the collagen bundles resemble the various features of TACS. We show these collagen bundle structures elicit cancer cell contact guidance, directional migration, and invasion differently from the traditional isotropic collagen nanofiber structures. This easy and low-cost fabrication technique of microscale collagen bundles may provide new possibilities of developing more physiologically relevant *in vitro* tumor models.

## 2. Materials and Methods

### 2.1. Collagen matrix preparations

Eight parts of type I bovine collagen monomer solution (3.0 mg/mL, pH 2, PureCol, Advanced BioMatrix, USA) were mixed with one part of 10× phosphate-buffered saline (PBS). The solution was then neutralized to a pH of ∼7.4 with about one part of 0.1M sodium hydroxide (NaOH) solution. Neutralization was conducted on ice. The collagen concentration of the neutralized solution was 2.4 mg/mL. In this study, we prepared all the samples with different collagen structures by mixing the same quantity of water or 1× PBS, to produce a final collagen concentration of 1.2 mg/mL.

For the ease of description, here we term the commonly used, traditional collagen gelling protocol (based on the manufacturer’s manual) as “Fiber method” and our novel collagen bundle fabricating protocol as “Bundle method” (Figure 1). For the fiber method, we thoroughly mixed 2.4 mg/mL neutralized cold collagen solution with the same amount of cold ultrapure water with a vortex mixer for ∼ 30 seconds. This diluted 1.2 mg/mL collagen precursor was then allowed to gel in a 37°C, humidified incubator. For the bundle method, instead of cold water, we vortex-mixed a neutralized cold collagen solution with the same volume of pre-warmed ultrapure water for 30 seconds. Visible collagen bundles appear instantly after the mixing procedure. More details on our collagen bundle fabrication method are presented in the Results section.

**Figure 1.**
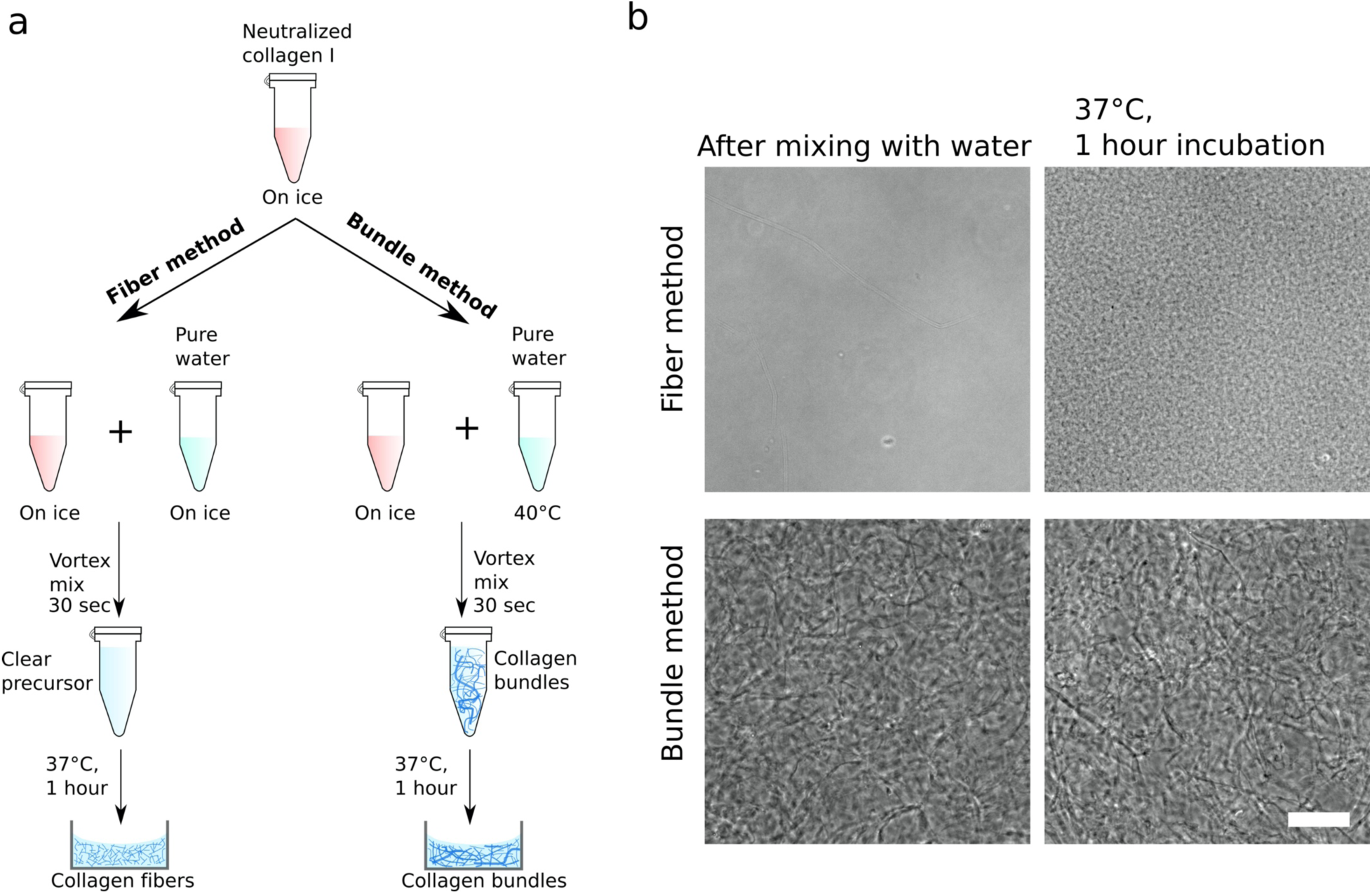
Rapid fabrication of collagen bundles comparing with collagen fibers. (a) The common procedure to prepare a collagen fiber gel (fiber method) versus the novel rapid fabrication of collagen bundles (bundle method). (b) Brightfield images comparing the appearance of the collagen-water solutions mixed using the fiber and bundle methods before and after incubation for 1-hour at 37°C. Scale bar: 100 μm. Using the fiber method, a clear solution of collagen and cold water polymerized into a collagen gel with dense fibrous structures after incubation. In the bundle method, visible bundles instantly formed after mixing with warm water and remained after the one-hour incubation.

### 2.2. Rat tail collagen extraction

Rat tail collagen was also used to demonstrate the collagen bundle fabricating technique. Sprague Dawley adult female rat tails were generously provided by Dr. Ryan Gilbert’s lab at RPI. Tails were initially frozen at -80 °C until processed. Tails were thawed at 4 °C overnight the day before extraction. For extraction, protocols similar to Ritte and Rajan *et al*. were used^27, 28^. Pliers were used to isolate the tail tendons, which were then submerged into a beaker of 1x PBS on ice. When all tails were processed, the tail tendons were rinsed three times with cold deionized water. Then, tail tendons were transferred to a new beaker containing 20 mM acetic acid, at a volume of 200 mL per tail. This beaker was placed on a magnetic stir plate with a stir bar at 4 °C and allowed to stir for 3 days. After, the solution was poured into 250 mL centrifuge bottles and centrifuged for 45 minutes at 10,000 ×g at 4 °C. The supernatant was collected into 50 mL centrifuge tubes and frozen at -80 °C overnight. The collagen solution was then lyophilized using a CentriVap (Labconco) and vacuum (Fisher) for 5 days. The resulting collagen mesh was collected, weighed, and resuspended at the concentration of 15 mg/mL in fresh 20 mM acetic acid in a glass vial and kept at 4 °C. To sterilize, chloroform was added at 10% of the volume and allowed to incubate overnight at 4 °C, where after the collagen solution was aseptically transferred into a sterile centrifuge tube. Purity of the collagen was verified via SDS-PAGE and mass spectroscopy by comparison of a commercially available rat tail collagen solution (Invitrogen).

### 2.3. Design and fabrication of the microfluidic chamber

We designed micro-post array patterns using the computer aided design (CAD) software SolidWorks (Dassault Systèmes), and the patterns were printed on a chrome mask by a high-resolution printing service (Front Range Photo Mask, CO, USA). A rectangular feature (thickness: 70-μm) with micro-wells in it was fabricated on a silicon wafer using a negative photoresist (SU-8 3050, MicroChem, MA, USA), through the techniques of photolithography. The standard fabricating guideline is provided by the SU-8 manufacturer. After the fabrication, we inspected the depth of the microwells by a stylus profilometer (Veeco, DekTak 8). We then molded the micro-post-containing rectangular chamber on the silicon master with PDMS (Sylgard 184, Dow Corning) following standard soft lithography techniques^27, 28^. After the PDMS was fully cured, we peeled it off and trimmed it to fit in a glass-bottomed cell culture dish. Before attaching the PDMS chamber onto the glass bottom, we treated the chamber with a plasma cleaner (Harrick Plasma) to make the PDMS surface hydrophilic. All the PDMS components were sterilized with 70% ethanol and then under UV light for at least 30 minutes.

### 2.4. Cell culture

In this study, we used an invasive breast cancer cell line MDA-MB-231, a human colon cancer cell line HCT-116, and a green fluorescent protein (GFP)-labeled MDA-MB-231 (a kind gift from Dr. Mihaela Skobe at Icahn School of Medicine at Mount Sinai). MDA-MB-231 and GFP-MDA-MB-231 cells were cultured in RPMI 1640 medium supplemented with 2 mM L-glutamine (Gibco), 10% FBS (Gibco), and 1% penicillin/streptomycin (Gibco). HCT-116 cells were cultured in McCoy’s 5A medium with 10% FBS and 1% penicillin/streptomycin. All cells were incubated in humid air maintained at 37°C and 5% CO_2_. The cell culture medium was changed every 2-3 days and the cells were passaged when they reached 90% confluency. For the 10-day cell culture in agarose-collagen co-gels, the cell culture medium was changed every three days.

### 2.5. Preparation of agarose and agarose-collagen co-gels

Low gelling temperature agarose powder (Sigma) was dissolved in ultrapure water or PBS in a 95°C water bath for 15 minutes. When fully dissolved, the water-based or PBS-based agarose solutions were kept in a 40°C water bath. The concentration (wt./vol.%) of the agarose solutions was 0.6%. In this study, we prepared three types of hydrogels: 0.3% pure agarose, 0.3% agarose-collagen fiber co-gel, and 0.3% agarose-collagen bundle co-gel. To make 0.3% agarose, we simply diluted a water-based 0.6% agarose solution with the same amount of pre-warmed PBS. To prepare 0.3% agarose-collagen fiber co-gel, the collagen solution was first neutralized on ice as described above. To minimize local gelation of agarose when it contacts the ice-cold neutralized collagen, the collagen solution was pre-warmed to 37°C for 5 minutes—during which no perceptible collagen polymerization occurred. We then thoroughly mixed the pre-warmed collagen solution with the same volume of PBS-based 0.6% agarose solution to produce an agarose concentration of 0.3%. For the 0.3% agarose-collagen bundle co-gel, we mixed the pre-warmed neutralized collagen solution with the same amount of water-based 0.6% agarose solution. In this way, the collagen content of both co-gels was 1.2 mg/mL. To grow cells in these gels, we quickly mixed a negligible volume of cell suspension at a cell density of 70,000 cells/mL in the gel precursors before they gelled.

### 2.6. Mechanical characterization of agarose and agarose-collagen co-gels

The mechanical characterization of the three gel types (pure agarose gels, agarose-collagen fiber co-gels, and agarose-collagen bundle co-gels) was performed on a high-precision piezo-electric actuator-controlled indentation system (CellScale, Canada). A small glass bead (radius *R* = 1.5 mm) attached on the end of a tungsten cantilever was used to indent the hydrogel samples. The indentation depth, *d*, was recorded by a camera and the indentation force, *F*, was computed based on the measured deflection and known bending rigidity of the cantilever. Using a MATLAB code, the experimental indentation force-depth curves were then fit into the Hertz contact model (Equation 1) for a rigid spherical indenter contacting a flat surface. The elastic modulus, *E*, of the hydrogel was calculated from this fit.

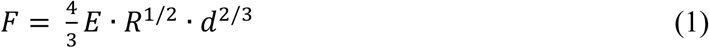

### 2.7. Immunofluorescence staining

The cell-containing hydrogel samples were washed in PBS, fixed with 4% paraformaldehyde at 37°C for 40 minutes, and permeabilized with 0.5% Triton X-100 at 37°C for 40 minutes. After washing with PBS three times for 30 minutes, the samples were blocked overnight in 5% BSA in PBS at room temperature. The samples were then incubated with rhodamine phalloidin (1:50, R415, Thermo Fisher) and Hoechst (0.5 μg/mL, Hoechst 33342, Thermo Fisher) protected from light and maintained at 4°C overnight.

### 2.8. Western blotting

To assess the expression of proteins associated with epithelial and mesenchymal phenotypes, Western blotting was performed. HCT116 and MDA-MB-231 whole cell lysates were isolated from flasks using mammalian protein extraction reagent (M-PER; ThermoFisher) following manufacturer protocols. Samples were treated with NuPAGE LDS Buffer (ThermoFisher) and NuPAGE reducing agent (ThermoFisher), and were heated to 70°C for 10 minutes. Protein separation was done through electrophoresis using a NuPAGE 3-8% tris-acetate gel (ThermoFisher) with wells loaded with 10 µg of protein. Subsequently, proteins were transferred onto a nitrocellulose membrane and then blocked overnight while shaken at 4°C in TBST-5% BSA – a TBS Tween-20 (TBST; ThermoFisher) solution containing 5% bovine serum albumin (BSA; Sigma).

Primary antibodies for vimentin (1:5000; Santa Cruz Biotechnology), E-cadherin (1:2000; Cell Signaling Technology), and GAPDH (1:15000; Cell Signaling Technology) were diluted in TBST-5% BSA. The diluted primary antibodies were applied to appropriated sections of the membrane overnight while shaken at 4°C. Membrane sections were washed overnight while shaken at 4°C in TBST. Horseradish peroxidase-conjugated secondary antibodies (Cell Signaling Technology) diluted in TBST-5% BSA were then applied for 1 hour. Secondary antibody dilutions were 1:10000 for vimentin, 1:2000 for E-cadherin, and 1: 15000 for GAPDH. Bound antibodies were detected using SuperSignal West Pico PLUS Chemiluminescent Substrate (ThermoFisher). Imaging was performed using a Chemidoc XRS+ system (BioRad). Quantification of band parameters were performed using the associated Image Lab software (BioRad).

### 2.9. Image acquisition

All bright field images of collagen fibers and bundles were obtained with an inverted microscope (Zeiss, Axio Vert.A1). Fluorescence images of cancer cells/tumor spheroids and reflectance images of collagen microstructures were acquired with laser scanning confocal microscopes (Leica SP8 or Zeiss LSM 510 META). Live confocal imaging of MDA-MB-231 seeded on 2D glass or embedded in collagen bundles or in collagen fibrous gels was conducted in an environmental chamber (Okolab) using a Leica SP8 for 5 hours at 15 min interval.

### 2.10. Scanning electron microscopy

Scanning electron microscopy (SEM) was used to characterize the microscopic structural differences between collagen fibers and bundles. The collagen fibrous gel prepared by Method 1 and the collagen bundles prepared by Method 2 were first fixed in glutaraldehyde at 4°C overnight. The samples were then dehydrated using a chemical drying method.^30^ In brief, after carefully rinsing the fixed samples with ultrapure water, we dehydrated the samples with a series of aqueous ethanol solutions: 50%, 75%, 80%, 90%, and 100%. The samples were allowed to stay in each ethanol concentration for 5 minutes and each ethanol concentration was applied twice. After the water in the samples was completely replaced by 100% ethanol, the samples were then plunged into hexamethyldisilazane (HMDS) for 40 minutes. Lastly, the HMDS was removed and the samples were air-dried. The dehydrated collagen fibers and bundles were sputter coated with platinum using a Denton Desk IV sputter coating system and imaged with a Zeiss SUPRA 55 FESEM in the Microscale and Nanoscale Cleanroom (MNCR) at Rensselaer Polytechnic institute.

### 2.11. Image analysis and statistical analysis

Length measurements from any brightfield images or confocal images were performed using ImageJ (NIH)^31^. Thicknesses of collagen fibers and bundles in confocal images were measured with the ImageJ plugin “Ridge Detection”^32^. Orientation of the microfluidics-aligned collagen bundles was characterized using the ImageJ plugin “OrientationJ”.^33^ The visualization and surface analysis software Imaris 9 (Oxford Instruments) was used to reconstruct confocal image stacks from which cell migration in 3D was automatically tracked and tumor volume was automatically computed. Data was presented as bar plots showing mean ± standard deviation (s.d), or as box plots, where boxes represent the 25th to 75th percentile with a median line, and whiskers represent the 1.5 interquartile range (IQR). Statistical difference between multiple experimental conditions was determined by one-way ANOVA tests with Tukey post hoc testing. Differences were considered significant at *p* < 0.05.

## 3. Results

### 3.1. Rapid fabrication of microscale collagen bundles

Toward the need for physiologically relevant *in vitro* ECM models, we present a procedure to rapidly fabricate microscale collagen bundles. The common method (termed “fiber method”, Figure 1a) of preparing collagen hydrogels includes three steps: (i) neutralization of the collagen monomeric stock solution (pH = ∼7.4, on ice, see details in Materials and Methods), (ii) adjustment of the collagen concentration by addition of ice-cold water, and (iii) gelation at 37°C for at least an hour. With this method, a thoroughly mixed, clear collagen precursor gradually polymerizes into a gel consisting of dense reticular collagen fibers that are on the order of hundreds of nanometers in thickness. In contrast, using the “bundle method” (Figure 1a), we found that replacing the ice-cold water in step (ii)—adjustment of collagen concentration—with warm water (40°C) resulted in visibly thick collagen bundles appearing instantly after mixing. The bundled collagen remained after an extended incubation at 37°C. Phase contrast microscope images (Figure 1b) show the appearance of collagen solutions at the different steps of the fiber and bundle methods, corresponding to Figure 1a. The drastic structural differences may be visualized in confocal reflectance images (Figure 2a), in which we also measured the thicknesses of the collagen fibers (mean ± s.d: 0.9 ± 0.5 μm) and bundles (mean ± s.d: 6.3 ± 1.7 μm) (Figure 2b). This method is not restricted to the bovine collagen utilized here, but functions the same using type I collagen isolated from other species (e.g., rat tail, Figure S2).

**Figure 2.**
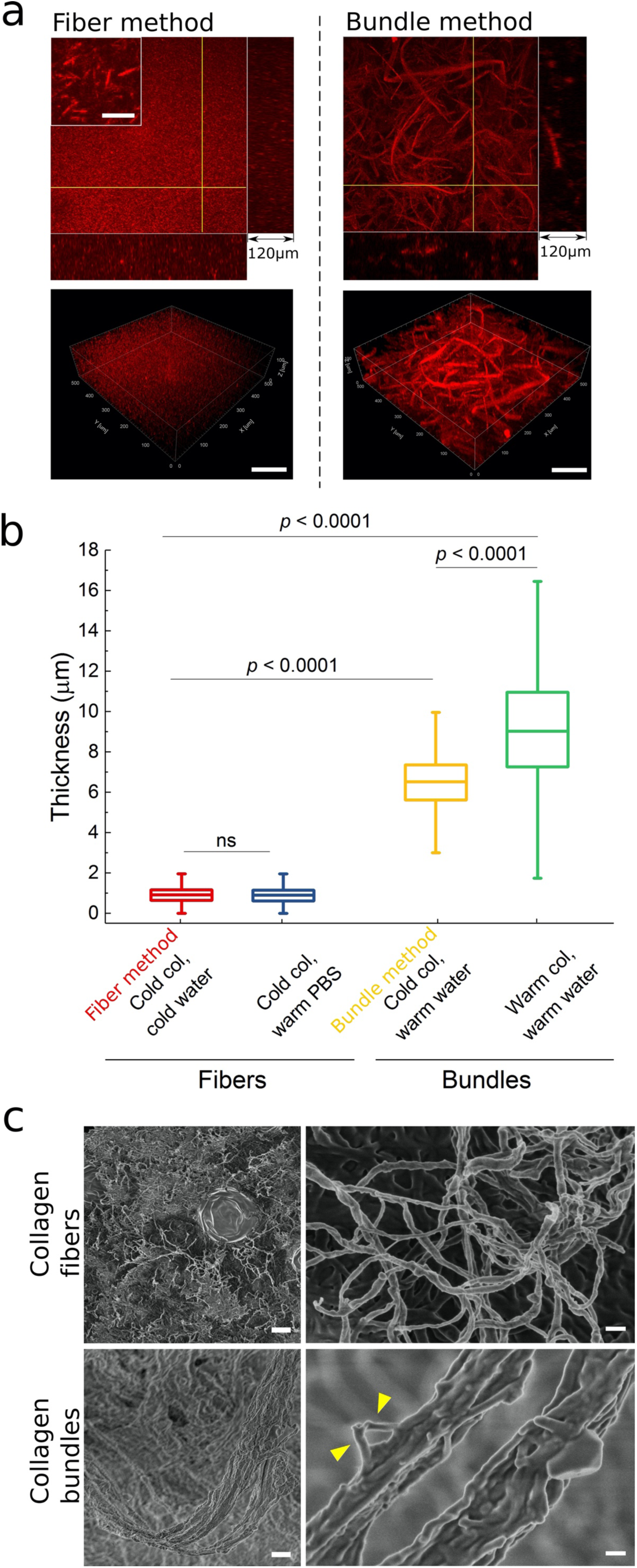
Morphological characterizations of collagen bundles compared to collagen fibers. (a) Confocal reflectance images of collagen fibers (scale bar in inset: 10 μm) and collagen bundles made by the respective methods. Scale bars in 3D reconstructions: 100 μm. (b) Characterizing the thickness of collagen fibers and bundles prepared by different protocols. The red box corresponds to the fiber method and the yellow box corresponds to the bundle method. One-way ANOVA with Tukey post hoc testing was performed. (n ≥ 1731 measurements, three samples each condition). Boxes represent 25th to 75th percentile and whiskers represent 1.5 IQR. (c) SEM images of microscopic structures of single collagen fibers (top row) and collagen bundles that consist of multiple bundled collagen fibers (bottom row). Scale bars: 2 μm (left column); 200 nm (right column). Arrowheads indicate free ends of collagen fibers in a collagen bundle.

Comparing the fiber and bundle methods, it is clear that introduction of *warm* water to the ice-cold neutralized collagen solution played a key role in the collagen bundle formation. To further investigate the bundling mechanism, we replaced the warm water with warm PBS (1×), which introduced a higher ionic strength due to the dissolved salts. Interestingly, mixing neutralized collagen with prewarmed PBS did not produce collagen bundles. Instead, the precursor turned into a gel with dense, reticular collagen fibers with thicknesses statistically similar to those produced by the fiber method (blue box, Figure 2b). Therefore, for collagen bundle formation, both increased temperature and low ionic strength of the diluting solution are necessary. The size of the bundles may be further manipulated by altering the temperature of the neutralized collagen precursor. A brief incubation at 37°C of the neutralized collagen precursor before mixing it with warm water significantly increased the thickness of the collagen bundles ((mean ± s.d: 9.0 ± 3.4 μm) green box, Figure 2b). The collagen morphologies corresponding to the four gelling conditions discussed in this section are shown in Figure S1.

To better understand the structure of the collagen bundles we produced, the samples were imaged with a scanning electron microscope (SEM, Figure 2c). When visualizing the microscopic structure, the much thicker collagen bundles were found to be composed of multiple long collagen fibers that coiled together, similar to *in vivo* observations^8, 25^.

### 3.2. Microfluidics-driven patterning and alignment of collagen bundles mimicking TACS-3

After fabrication, the collagen bundles were randomly and loosely packed in solution and did not form a hydrogel like the fiber method. Since they were not bound to a hydrogel, these micro-scale collagen bundles could be used to form large-scale, complex patterns with controllable curvature and local alignment. The collagen bundle alignment resembles the aligned, bundled collagen of TACS-3. We aligned the bundles into different micro-scale curvatures using a post-production method based on microfluidics (Figure 3). A PDMS microfluidic chamber (width: 15 mm, height: 70 μm) with arrays of cylindrical (diameter: 70 μm) and triangular (equilateral, side length: 830 μm) micro-posts was made. The chamber had an open end where the collagen bundles were introduced, and tubing was sealed to a port on the opposite end so that a vacuum could be applied across the chamber (Figure 3a). A syringe was attached to the free end of the tubing and used to apply a gentle vacuum to pull a solution containing the collagen bundles through the chamber. The long collagen bundles were captured by the micro-posts and the flow facilitated the alignment of the free ends of these bundles (Figure 3b). The combination of the micro-posts’ geometric features and the direction of the flow arranged the collagen bundles into unique patterns (Figure 3c). When observing the collagen bundles around a micro-post, we found that the bundles were locally aligned, and the orientation of their local alignment was determined by the cross-sectional geometry of the micro-post (Figure 3d).

**Figure 3.**
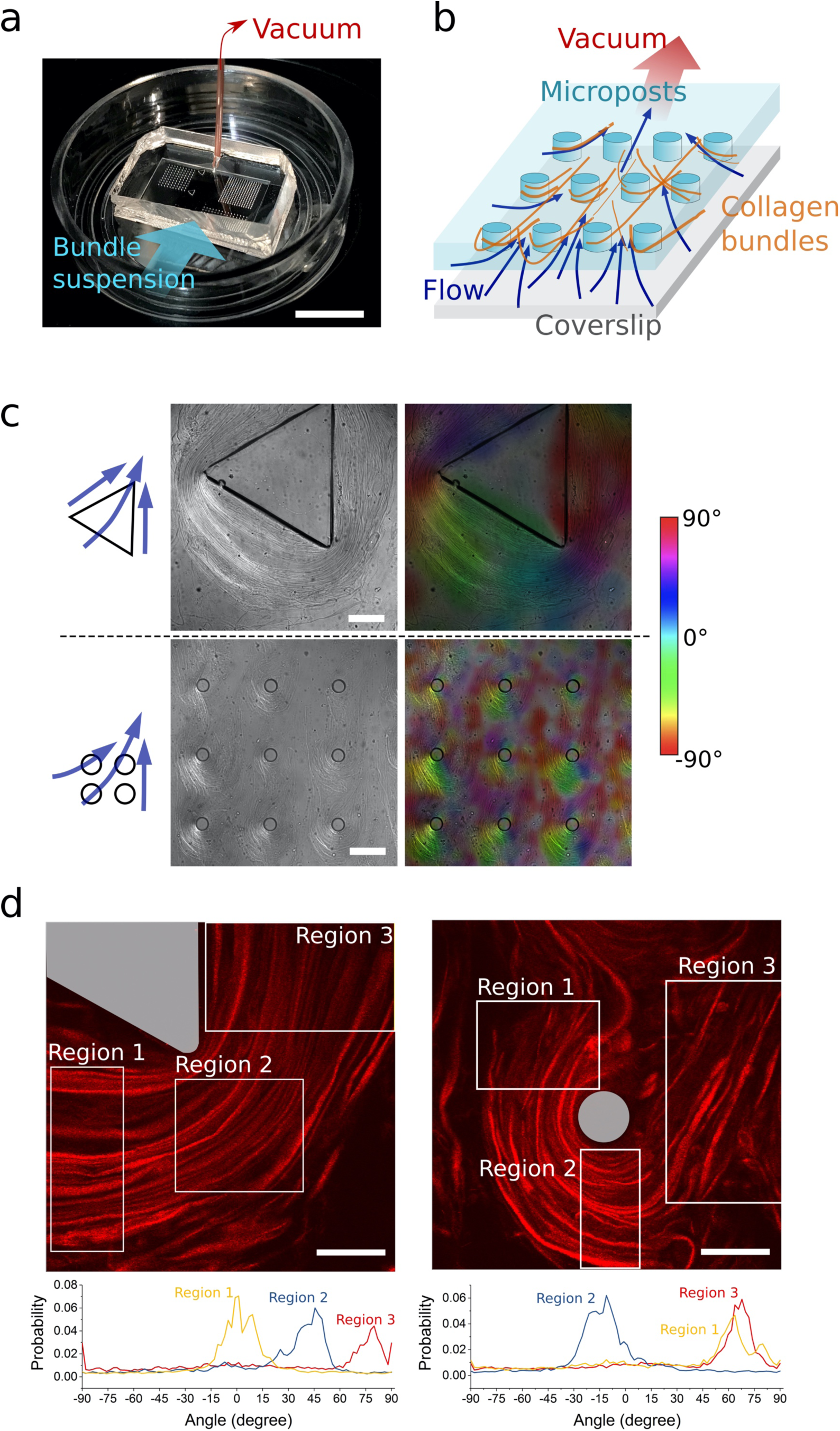
Microfluidics-driven patterning and alignment of collagen bundles. (a) Microfluidic chamber containing micro-post arrays. A suspension of collagen bundles was introduced at the open end and a vacuum was applied through the tubing sealed to the opposite end. The vacuum pulled the suspension through the micro-chamber. Scale bar: 10 mm. (b) Schematic showing the design of the microfluidic chamber and mechanism of microfluidic flow driving collagen bundles to pass through and align around an array of micro-posts. (c) Large-scale differential interference contrast (DIC) images of collagen bundles near micro-posts of varying shapes (left) and their respective colormaps that denote bundle orientation (right). Blue arrows indicate the flow direction corresponding to the DIC images. Scale bars: 200 μm. (d) Confocal images of collagen bundles around micro-posts of different shapes. Cross-sections of the micro-posts are highlighted in grey. Below the confocal images we show the orientation characterizations of the bundles in multiple regions in each image. Scale bars: 100 μm.

### 3.3. Collagen bundle arrangement facilitates cell contact guidance

The patterned microscale collagen bundles resembled the alignment of thick collagen bundles observed in *in vivo* studies^10, 25^, specifically TACS-like aligned collagen bundles. To investigate the interactions between cancer cells and the patterned collagen bundles, we flowed metastatic breast cancer cells MDA-MB-231 (GFP-labeled) with the collagen bundles through the microfluidic chamber and cultured them *in situ* for one week. Not only did the cancer cells grow along the aligned collagen bundles with the same curvature (Figure 4a), but upon closer inspection it can be seen that the cancer cells inserted themselves in between the collagen bundles (Figure 4b, c). When caught by the micro-posts from the flow, the bundles distributed themselves over the entire height of the micro-posts creating arrangements of bundles multiple layers in height as well as thickness (radially from the center of the post). The cancer cells were able to migrate between the collagen bundles (Figure 4d) because the spacing between the aligned collagen bundles was on the order of a few tens of micrometers, which is approximately the size of a single cell (Figure 4e). With this assay we showed that the 3D aligned collagen bundles enabled cancer cell contact guidance.

**Figure 4.**
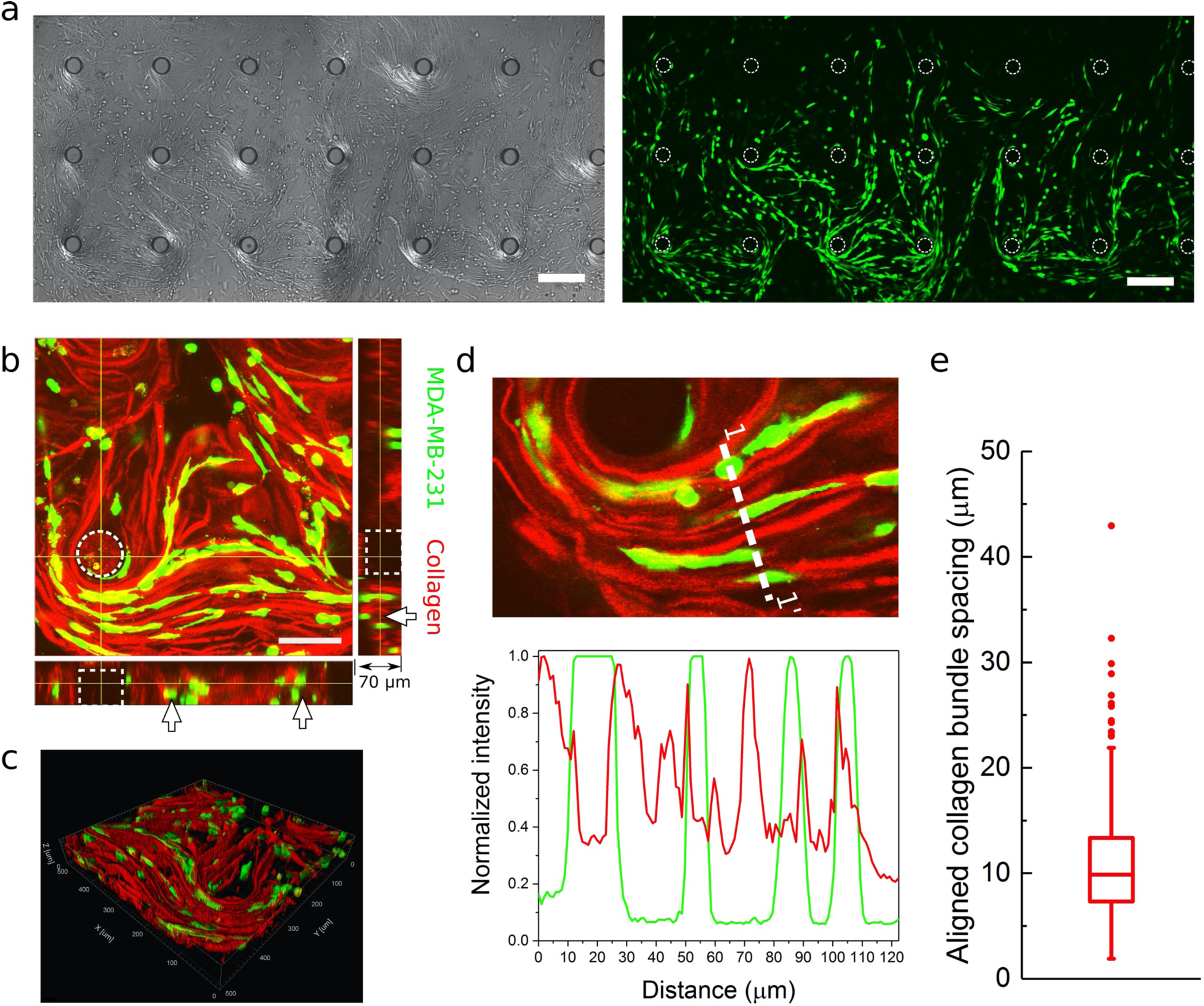
Complex collagen bundle topography facilitates cell contact guidance. (a) DIC images (left) and confocal images (right) showing large scale collagen bundle topographic landscape guiding directional growth of GFP-labeled MDA-MB-231 cells. Scale bars: 200 μm. (b) Confocal image volume showing cells reoriented to the local collagen bundle structures. Arrows indicate the cells and collagen bundles were distributed through the height of the microfluidic chamber. (c) 3D reconstruction of cell contact guidance in collagen bundle topography. (d) Spatial characterization showing that cancer cells insert themselves in between the collagen bundles, mimicking *in vivo* observations. (e) Spacings between collagen bundles measured from confocal reflectance images (n = 332 measurements). Individual data points are outliers.

### 3.4. Aligned collagen bundles enhance cell motility and directional migration

After establishing cancer cell contact guidance in the 3D aligned collagen bundles, we characterized the migration of the cancer cells in the 3D aligned collagen bundles compared to both 2D collagen-coated glass and 3D reticular dense collagen fibers. After 24-hours of culture, we imaged MDA-MB-231 cells in the three conditions (Figure 5a) every 15 minutes for five hours using confocal 3D live imaging (Figure 5b, Movie S1). Both migration speed and persistence of the cells in the three conditions were characterized.

**Figure 5.**
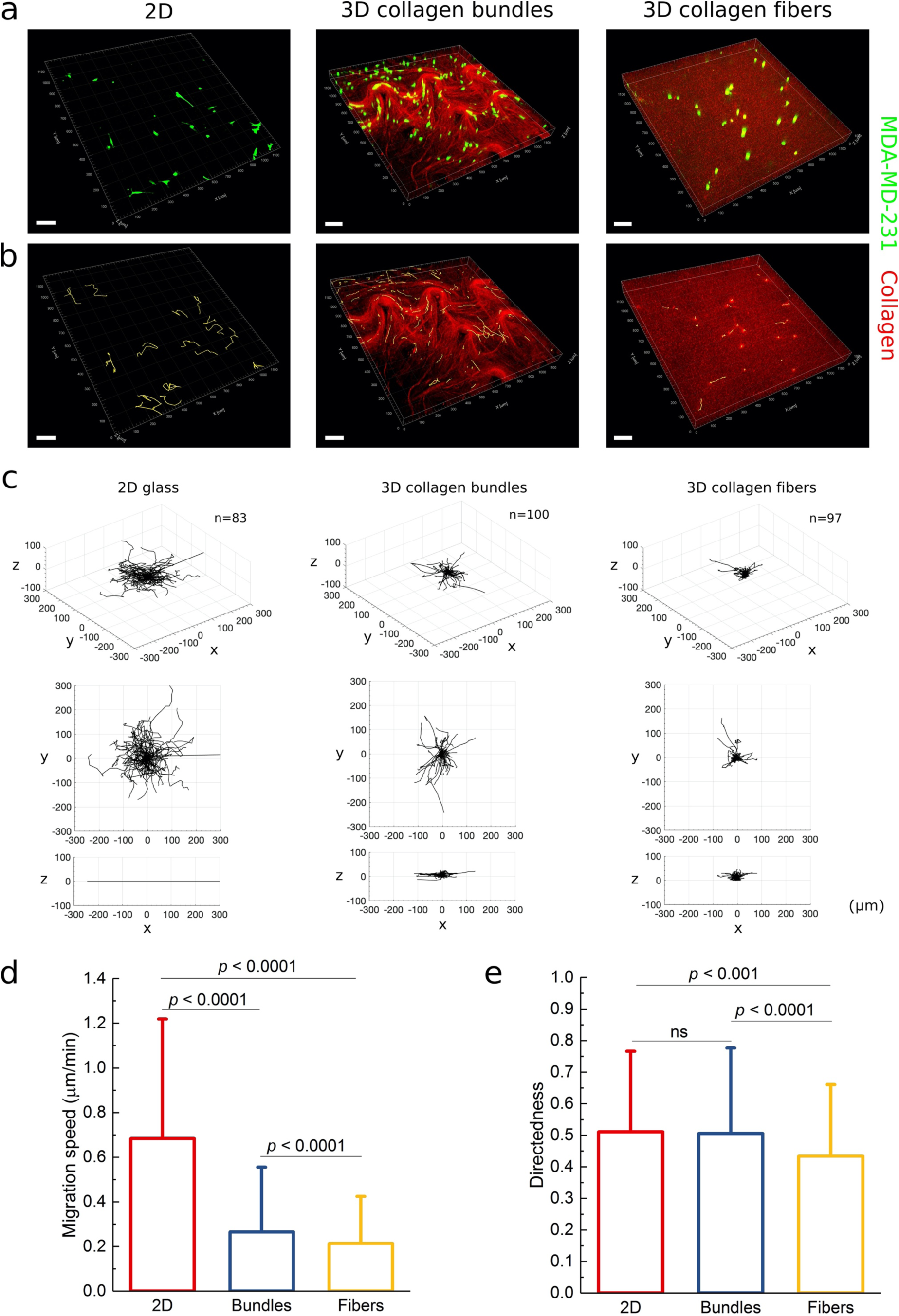
Cancer cell migration characterization under varied collagen structural conditions. (a) Confocal images of MDA-MB-231 cells seeded on collagen coated glass, in between collagen bundles, and in dense reticular collagen nanofibers. (b) Migration tracks of MDA-MB-231 cells under the three culture conditions over five hours. Yellow lines denote the cell migration trajectories. Scale bars: 100 μm. (c) Three-dimensional cell trajectory plots comparing cell migration patterns under the three collagen-structural conditions. (d) Frame-to-frame cell migration speeds between 15-minute time interval (n ≥ 3323 measurements of at least 206 cells) and (e) directedness of the cells over a five-hour period (n ≥ 206 cells).

To compare migration speeds, we found the average frame-to-frame speed (the displacement of a cell between two frames divided by the time interval, 15 minutes) for each of the three conditions (Figure 5d, Movie S2). Expectedly, without constraint in the third dimension the cells on the 2D glass surface were most mobile; they moved fastest at a mean speed of 0.68 μm/min. The cells in both collagen conditions were significantly slower than on 2D glass. The mean speed of the cells between aligned collagen bundles (0.27 μm/min) was the second fastest, which, although close to, was significantly faster than that of the cells embedded in dense collagen fibers (0.21 μm/min). To compare the persistence of cells migrating in the three conditions, we used the cell directedness, defined as the ratio of cell displacement to the trajectory length^34^ (also see Figure S3, Figure 5e). The cell directedness measurement is lowest for the cells embedded in the collagen fibers, which indicates they were least persistent. The cells on 2D glass and in the collagen bundles had comparable directedness values.

The speed and persistence quantifications are corroborated by visualizations of *in situ* cell trajectories and three-dimensional wind rose plots. Five-hour long trajectories are superimposed in yellow onto the respective collagen structures in Figure 5b. The cells on 2D collagen-coated glass migrated the furthest and their trajectories were random while the trajectories of most cells in the collagen bundles matched the local bundle topography. In comparison, the cells embedded in collagen fibers were less migratory and their trajectories extended least into the collagen matrix. Visualizing these trajectories as 3D wind rose plots confirms the 2D, relatively long migration paths of the cells on glass and the more directed motion of the cells migrating on the bundles compared to the fiber collagen gel (Figure 5c). These results indicate that aligned collagen bundles enhance the speed and the translocating capability of the cells over a fiber collagen gel.

### 3.5. Agarose-collagen bundle co-gels as a novel platform for recapitulating carcinoma cell *in vivo* phenotypes

Having shown that collagen structure and dimensionality influenced cancer cell motility, we then investigated whether collagen structures play a role in model tumor growth. We chose to grow the model tumors in collagen-agarose co-gels since they allow the mechanical stiffness to be decoupled from collagen density.^35, 36^ Specifically, three types of hydrogels/co-gels were prepared: (i) pure agarose, (ii) agarose-collagen dense fiber co-gel, and (iii) agarose-collagen bundle co-gel (Figure 6a). Both the agarose and collagen content were kept constant across all three gel types, at 0.3% (wt./vol.) and 1.2 mg/ml, respectively. The stiffness of the matrices is determined by the agarose content. Indeed, the three gels had similar mean elastic moduli values in the range of 267 Pa – 336 Pa, which is similar to that of healthy mammary tissue^37^, (Figure 6b). Although the collagen content (and therefore ligand density) was also kept constant, the difference between the co-gels is the collage structure: in one the dense fiber collagen was incorporated and in the other collagen bundles were incorporated. In Figure 6a visual comparisons of the gel structures may be seen at different length scales.

**Figure 6.**
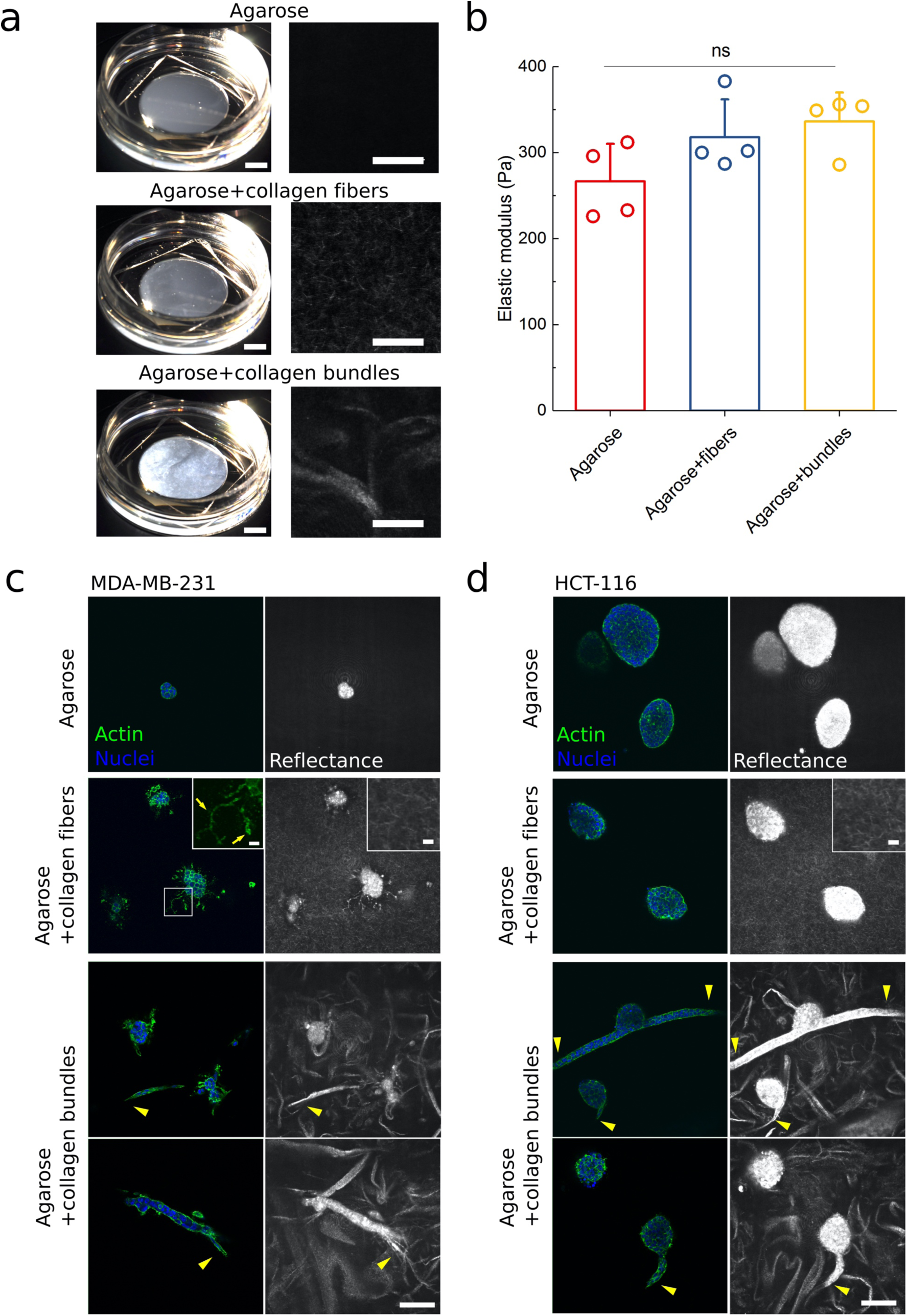
Agarose-collagen based 3D composite hydrogels as *in vitro* tumor models with tunable collagen structures. (a) Macrographs (left column, scale bars: 5 mm) and confocal images (right column, scale bars: 50 μm) of three types of gels: 0.3% (w/v) agarose, 0.3% agarose+1.2mg/mL collagen fiber co-gel, 0.3% agarose+1.2 mg/mL collagen bundle co-gel. (b) Elastic modulus characterizations of the three gel types. No significant difference was detected in the elastic moduli of these gels. (One-way ANOVA with Tukey post hoc testing, n = 4 for each gel type). (c) MDA-MB-231 tumor growth (Day 10) in the three gel types. Scale bar: 100 μm. Insets show the filopodia (yellow arrows) extending into the surrounding agarose-collagen fiber matrix. Scale bars in insets are 10 μm. (d) HCT-116 tumor growth (Day 10) in the three gel types. Scale bar: 100 μm. All the arrowheads indicate the multicellular invasive growth along collagen bundles.

Breast cancer cells (MDA-MB-231) and colon cancer cells (HCT-116) were cultured in the three gel conditions. As shown in literature^38, 39^, these two cell lines of epithelial origin have different metastatic potentials: the MDA-MB-231 cell line is known to have a higher metastatic potential, whereas the HCT-116 cell line retains more epithelial characteristics (also see expression of epithelial-mesenchymal transition markers in Figure S4). After 10 days of growth in the matrices the cells were fixed, and their F-actin and nuclei were stained so that their interactions with the matrices could be visualized.

The growth behavior and morphologies were different both between the different types of cancer cell and between the gel types. Pure agarose provided the cells with 3D mechanical constraint without any adhesive ligands. Both the MDA-MB-231 and the HCT-116 cells grew into tumors with well-defined, smooth boundaries (Figures 6a and 6b, top row). However, the HCT-116 cells more readily proliferated in this environment to form larger masses (Figure 6b, top row). The presence of dense collagen nanofibers in the gels induced more proliferation from the MDA-MB-231 cells creating larger and more irregular mass shapes, which were punctuated by many long extended filopodia into the surrounding matrix showing a tendency of invasion (Figure 6a, second row). Interestingly, the addition of dense collagen nanofibers to the agarose gel did not have a significant effect on the mass size or morphology of HCT-116 cells (Figure 6b, second row). Large-scale confocal image volumes (Figure 7a) facilitated the measurement of tumor volume, which was quantified for each cell and gel type and is presented in Figure 7b.

**Figure 7.**
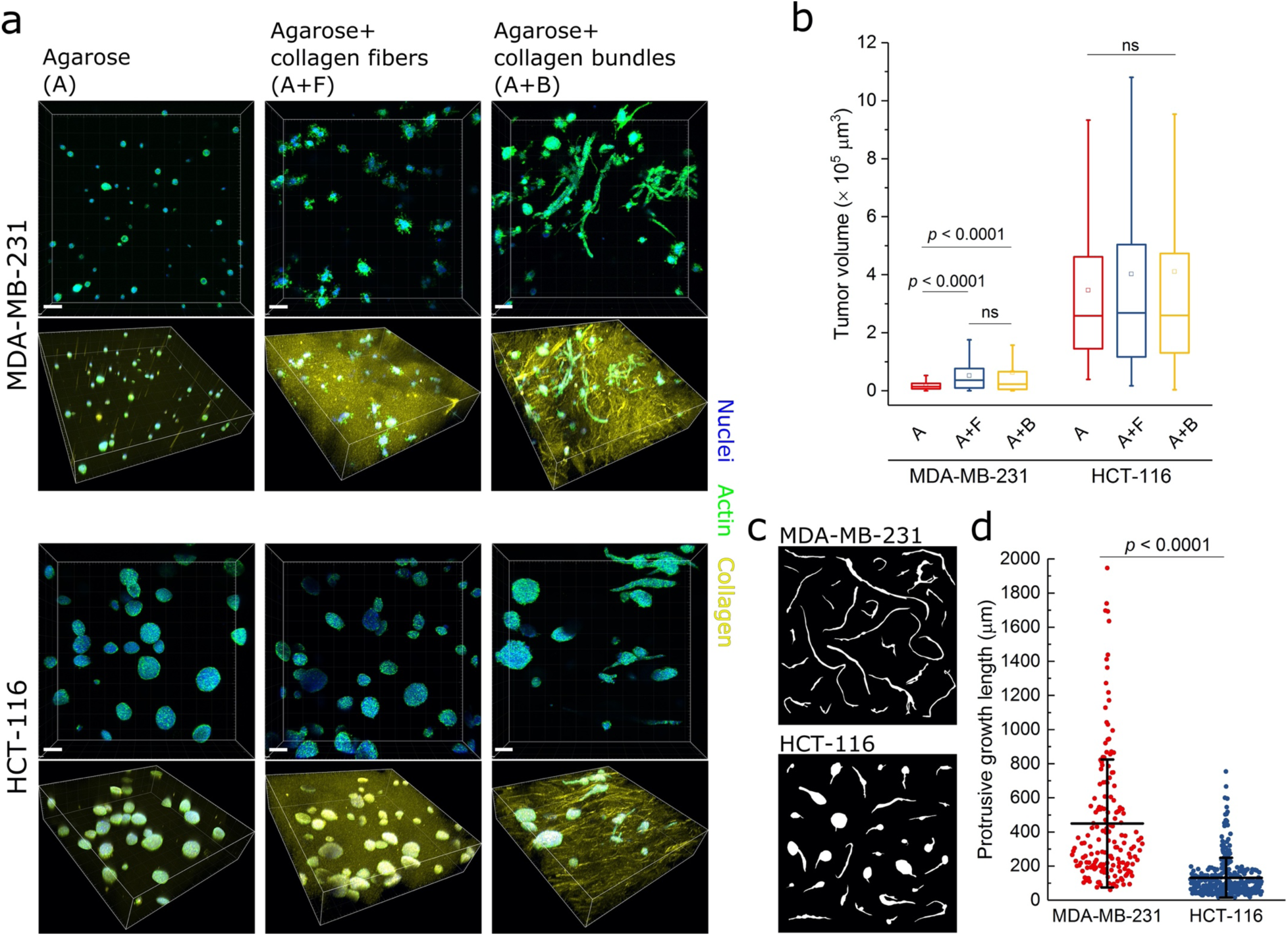
Collagen bundles in agarose mimic TACS-1 and elicit cancer invasion. (a) Large scale 3D confocal imaging of MDA-MB-231 and HCT-116 tumors cultured for 10 days in pure agarose (A), agarose-collagen fiber co-gel (A+F), and agarose-collagen bundle co-gel (A+B). Scale bars: 100 μm. (b) Box plot showing tumor volume measurements of the two cell lines in the three gel conditions. One-way ANOVA with Tukey post hoc testing was used to compared three gel condition within each cell line. (n ≥ 326 tumors for HCT-116; n ≥ 621 tumors for MDA-MB-231) (c) Morphology of protrusive growth of both cell lines elicited by collagen bundles in co-gels. (d) Protrusive growth length measurements of both cell lines (mean ± s.d). One-way ANOVA with Tukey post hoc testing, n ≥ 168 protrusive growths, two samples each condition.

When cultured in agarose-collagen bundle co-gels, MDA-MB-231 cells formed two types of morphology depending on their location with respect to a collagen bundle. In the case that they grew well away from a collagen bundle, their morphology was the same as in the collagen-fiber gels. If they grew adjacent to a collagen bundle, the cells invaded and grew along the bundle as a cohort and formed long strands (mean ± s.d.: 450 ± 375 μm) such that masses outside of the bundles were not present (Figure 7c). The same observations held for the HCT-116 cells, however, to a lesser extent. The HCT-116 cells tended to maintain tumor masses outside of the bundles— growing against the agarose matrix—with significantly shorter cell cohort protrusions (mean ± s.d.: 132 ± 116 μm) into the collagen bundles. This might be explained by the retention of the epithelial phenotype by the HCT-116 cells and the strong cell-cell junction strength due to high E-cadherin expression.

Through the observations of the growth behaviors and morphologies of the two cancer cell lines we learned that invasive behaviors of cancer cells do not only require the presence of collagen, but also particular dimensions and structures of the collagen component provided by the collagen bundles. Furthermore, the MDA-MB-231 breast cancer cell line is known to be more aggressive and metastatic than the HCT-116 colon cancer cell line. The phenotypes of the two cell lines are recapitulated in our co-gels where the HCT-116 cells generally retained a more epithelial phenotype and corresponding markers (Figure S4). Combining our microfluidic-drive alignment and the co-gel production, we were also able to demonstrate an agarose-collagen co-gel with complex collagen bundle patterns and the capability of repeatedly observing the tumor growth at the same locations in the co-gel (Figure S5).

## 4. Discussion

In this study, we show a simple, rapid, and robust method of fabricating micro-scale, type I collagen bundles, which are not achievable using a traditional collagen gelling protocol. We showed that this method is applicable to type I collagen isolated from different species (bovine skin and rat tail). By varying temperature and ion content in the collagen precursor, we found high temperature and low ionic strength played key roles in the instant collagen bundle formation. As widely shown in *in vivo* studies, the tumor microenvironment often contains abundant microscale collagen fiber bundles that could directly guide cancer cell invasion as single cells or in a collective way^10^. This biological process may not be fully recapitulated by traditional collagen fiber gels due to the lack of the proper collagen architecture and dimension. Attempting to fill this gap, we evaluated how cancer cells interacted with our novel collagen bundles compared to traditional collagen nanofibers.

Making use of microfabrication and microfluidic techniques, we first demonstrated a novel and flexible way to post-process the collagen bundles into large scale controllable complex patterns with local alignment that mimics TACS-3 – the aligned thick collagen strands that facilitate cell invasion. This complexity may not be created using other techniques, such as straight microfluidic channels^22^ or electrospinning^40^. In *in vitro* collagen nanofiber models, it is well known that local collagen remodeling by cancer cells enhances cell invasion^7, 41^. Our results showed that dense collagen nanofiber meshes trapped the cancer cells and limited cell motility before they were able to significantly remodel the collagen. Due to cell-matrix adhesion, cell migration speed was slowed down in both 3D collagen compared to the 2D case, however, the cell-scale microtracks provided by the microfluidics-aligned collagen bundles significantly enhanced cancer cells’ contact guidance and their directional migration.

We then demonstrated the production of an interpenetrating network hydrogel consisting of collagen bundles and agarose, different from an early innovation of an agarose-collagen fiber co-gel^35^. Although the previous co-gels indeed provided new perspectives of creating physiologically relevant models with a controllable stiffness and collagen density, they did not emphasize the importance of the architectural factors. By controlling gelling temperature and ion content, we produced an agarose-collagen bundle co-gel and an agarose-collagen fiber co-gel with comparable stiffness at the same agarose concentration.

We compared pure agarose, agarose-collagen bundle co-gels, and agarose-collagen fiber co-gels as *in vitro* 3D tumor models. The growth of two cancer cell types with different metastatic potentials responded to the three gel types differently. Under the same mechanical stiffness, the presence of collagen and the collagen structures drastically influenced the behavior of invasive breast cancer cell MDA-MB-231. Although the presence of collagen nanofibers significantly enhanced the cell proliferation and elicited an invasive tendency of the mechanically constrained cells, cell invasion could not be achieved. On the other hand, in the collagen bundle-containing agarose, cancer cells underwent collective migration along the collagen that has been seen *in vivo*. In comparison with MDA-MB-231 cells, human colon cancer cells HCT-116, with strong cell-cell adhesion, did not appear to have a strong invasive tendency when surrounded by collagen nanofibers. However, in the agarose-collagen bundle co-gel, distinct cell-cohort protrusions originating from HCT-116 tumor spheroids were observed when the growing tumors made contact with the collagen bundles.

Traditional methods of making nanoscale collagen fibers with variable fiber thickness have been well-established and are reproducible. With the characterizations of cancer cell motility and invading capability, our novel yet simple method of rapidly producing microscale collagen bundles undoubtedly pushes forward the controllability of collagen structure as *in vitro* tumor models. One may use the collagen bundle-based *in vitro* models to better understand the underlying mechanism of cell invasion along TACS and in fibrotic stroma, such as energy cost or the role of metastasis-responsible proteins, for better translational outcomes. Not only may the collagen bundles be applied to study cancer cell phenotypes and metastasis, but also provide new avenues for tissue engineering using a natural tissue component – collagen – as scaffolds. For example, aligned microscale bundled collagen may provide biocompatibility, adhesion, and proper topography for skeletal muscle growth^42, 43^ and nerve regeneration^44, 45^.

## 5. Conclusion

In summary, we have developed a novel and simple fabrication method to rapidly produce microscale collagen bundles, of which the thickness is comparable to TACS fiber bundles. These collagen bundles could be aligned by a microfluidic device or combined with agarose hydrogel to form an interpenetrating network co-gel. The TACS-3 mimicking aligned collagen bundles were shown to elicit cancer cells’ contact guidance and enhance their directional migration. In agarose-collagen co-gels, we showed two different cancer cells responded to collagen structures differently, and found that microscale collagen structures were required for both cell types to invade. Enabling cancer cell invasion has been determined to be one of the key features of TACS. Overall, our experiments using microscale collagen bundles elucidated the significant role of stromal collagen structure in cancer cell behavior. This study has expanded the controllability of reconstituted collagen structure, and therefore provided a possibility of creating more physiologically relevant *in vitro* 3D models.

## Acknowledgements

We thank Jamie Gearhart in the Mills Lab for the help with the mechanical testing on the hydrogels. We thank Prof. Lee Ligon for the helpful discussion. We acknowledge Dr. Sergey Pryshchep’s assistance on confocal microscopy, the National Science Foundation (Grant # NSF-MRI-1725984), and the software access (Imaris) in the Biotech Microscopy Core at Rensselaer Polytechnic Institute (RPI). We also thank M. David Frey for his technical assistance with scanning electron microscopy performed in the Micro and Nano Fabrication Clean Room (MNCR) at RPI. This work is supported by start-up funding for Dr. Mills’s lab from RPI.

