## Supplementary data for "Rapid fabrication of collagen bundles mimicking tumor-associated collagen signatures"

### Supplementary Information

Xiangyu Gong[1, 3], Jonathan Kulwatno[1, 2], K. L. Mills[1, 3, *]

[1]Department of Mechanical, Aerospace, and Nuclear Engineering, Rensselaer Polytechnic Institute, 110 8th St, Troy, NY 12180

[2]Department of Biomedical Engineering, Rensselaer Polytechnic Institute, 110 8th St, Troy, NY 12180

[3]Center for Biotechnology and Interdisciplinary Studies, Rensselaer Polytechnic Institute, 110 8th St, Troy, NY 12180

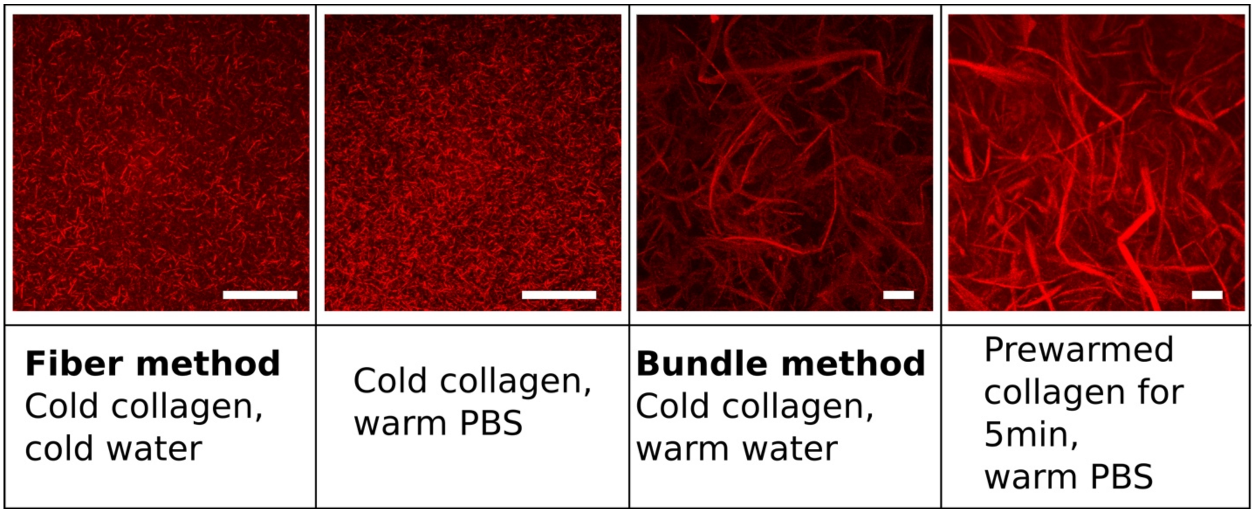


**Figure S1.** Temperature and ionic strength control collagen bundling. Confocal images show the morphologies of the collagen fibers (max projection of 47 μm) or bundles (max projection of 120 μm) prepared by the listed four different methods, corresponding to the thickness characterization in Figure 2b. Scale bars: Scale bars: 50 μm.

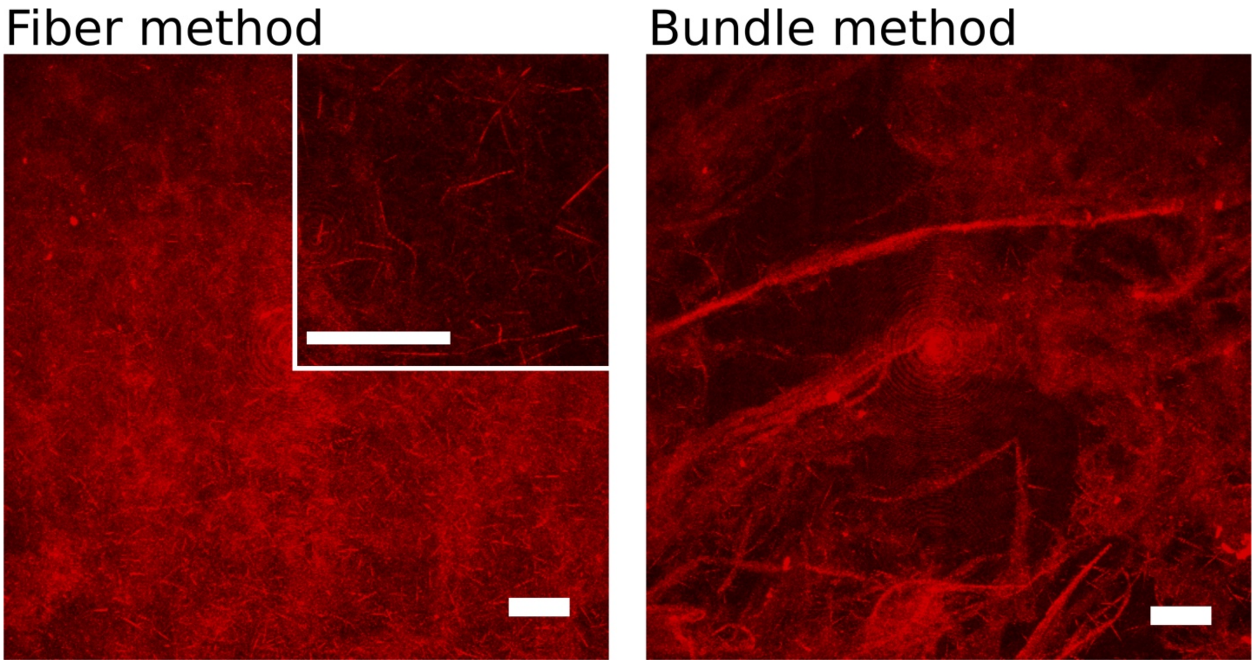


**Figure S2.** Collagen fibers and collagen bundles made of rat tail collagen type I (1.2 mg/mL) using the “fiber method” and the “bundle method”, respectively. Scale bars: 50 μm.

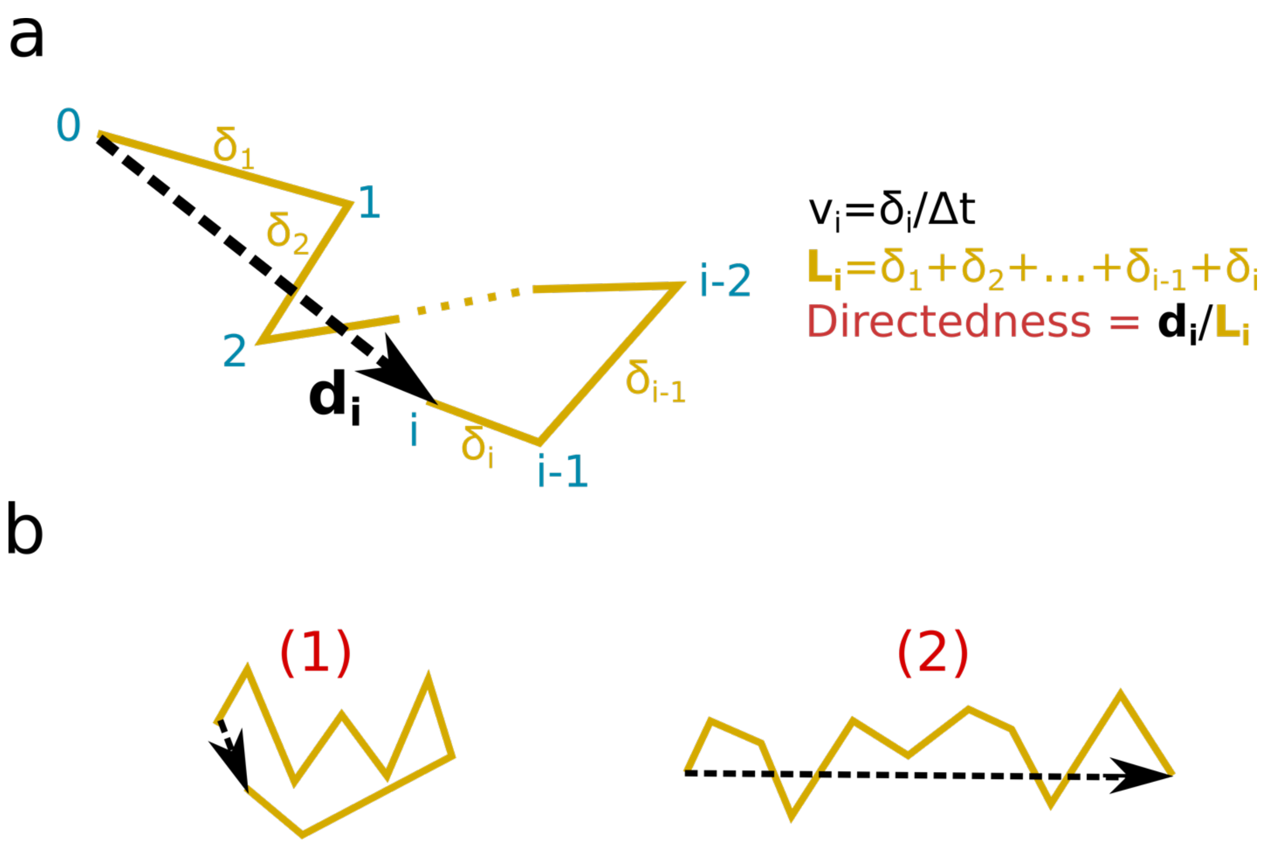


**Figure S3.** Cell migration speed and directedness. (a) Schematic of a cell trajectory recorded at a rate of Δt/frame, defining cell speed and directedness. (b) Example showing Trajectory (1) is less directed than Trajectory (2).

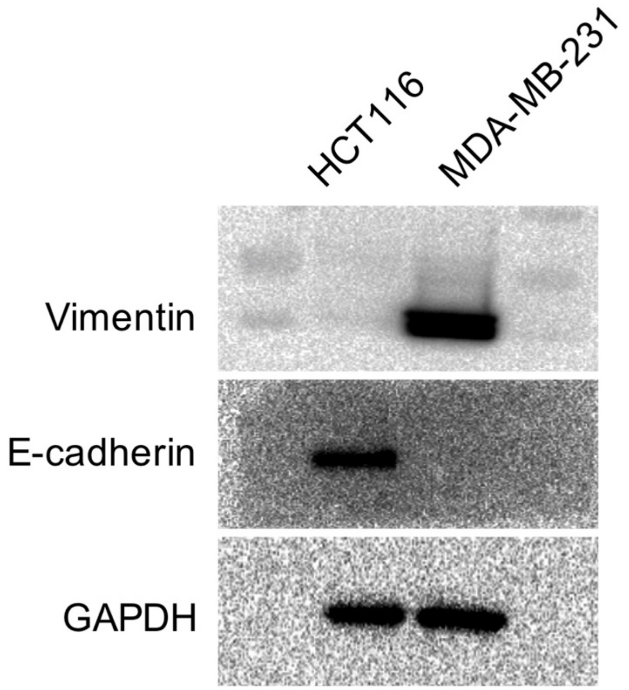


**Figure S4.** Protein expression of Vimentin and E-cadherin compares EMT protentional of human colon cancer cells HCT-116 and breast cancer cells MDA-MB-231.

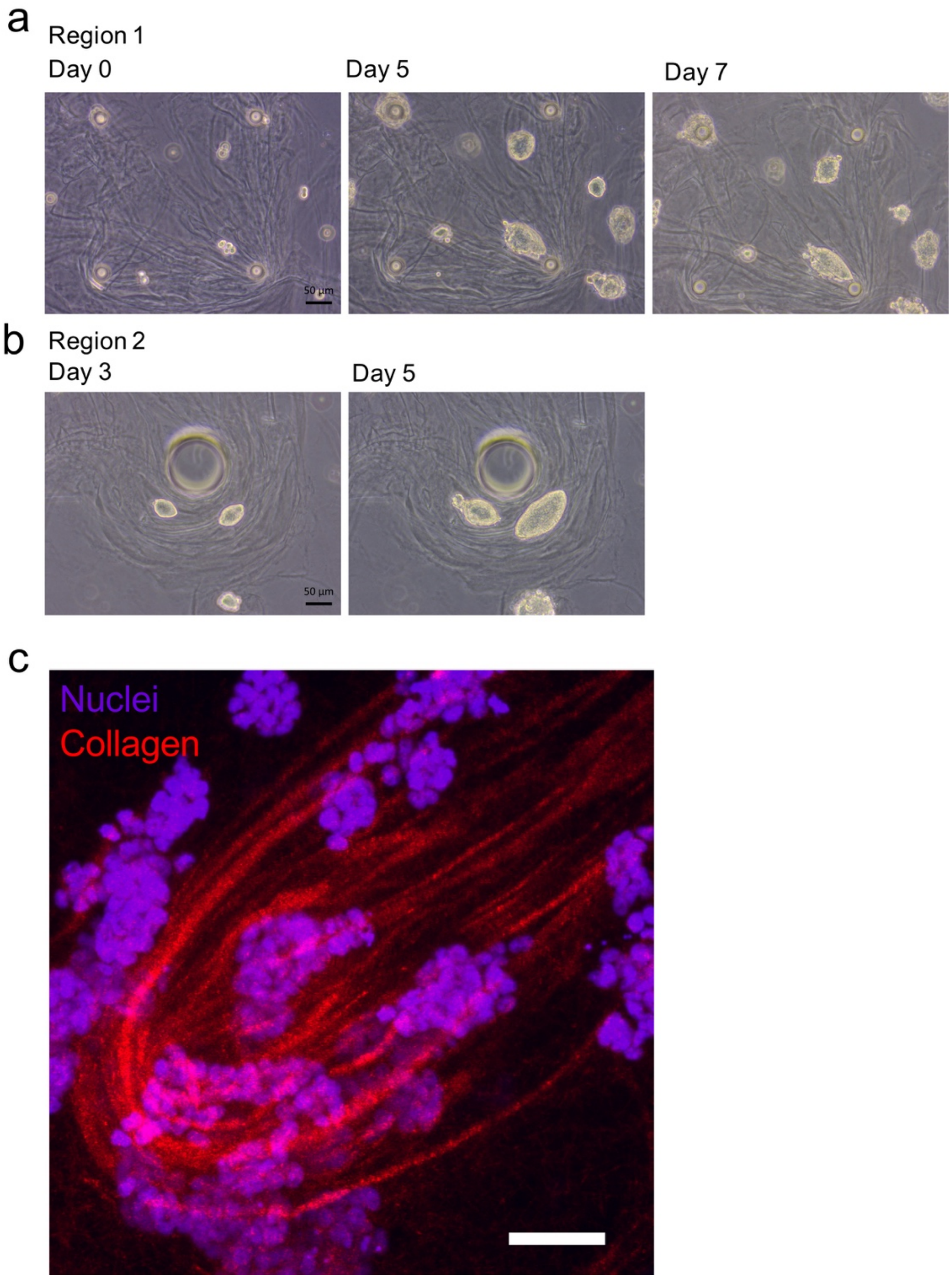


**Figure S5.** Microfluidics-driven collagen bundles in collagen-agarose co-gels. The agarose concentration is 0.3% (w/v). (a, b) Brightfield images tracking HCT-116 cells growing at two regions over 5-7 days in agarose containing aligned collagen bundles. (c) Confocal imaging showing HCT-116 multicellular tumors growing in an aligned collagen bundle-agarose cogel. Scale bar: 50 μm.

**Supplementary Movies**

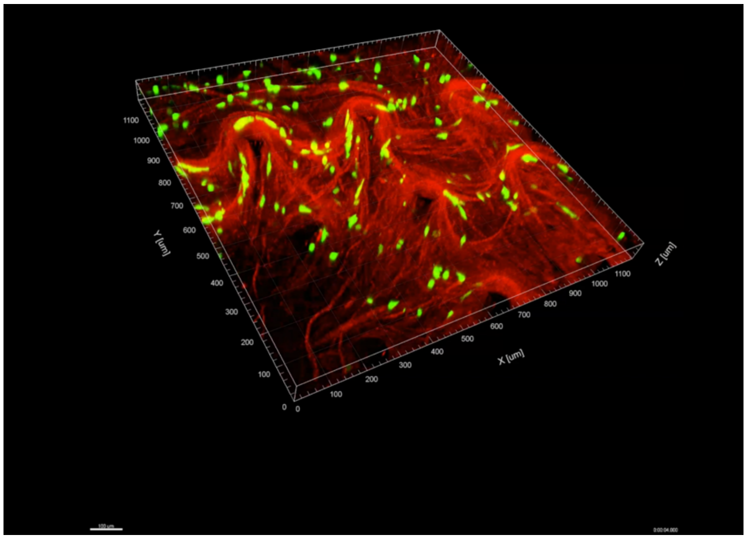


**Movie S1.** Cancer cells (MDA-MB-231) migrating under varied collagen structural conditions for five hours (green: cells; red: collagen).

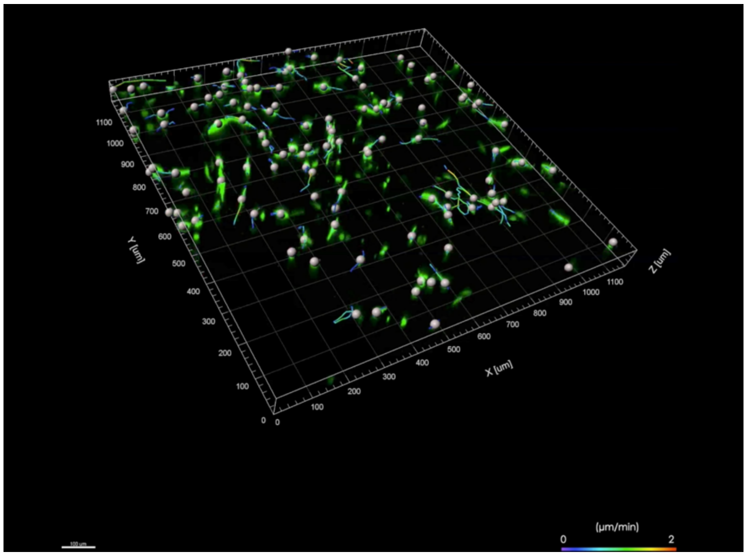


**Movie S2.** Trajectory of cancer cells (MDA-MB-231) migrating under varied collagen structural conditions for five hours. The trajectories are color-coded based on the real-time cell migration speed. Collagen is not shown in this movie.
